# Construction of thousands of single cell genome sequencing libraries using combinatorial indexing

**DOI:** 10.1101/065482

**Authors:** Sarah A. Vitak, Kristof A. Torkenczy, Jimi L. Rosenkrantz, Andrew J. Fields, Lena Christiansen, Melissa H. Wong, Lucia Carbone, Frank J. Steemers, Andrew Adey

## Abstract

Single cell genome sequencing has proven to be a valuable tool for the detection of somatic variation, particularly in the context of tumor evolution and neuronal heterogeneity. Current technologies suffer from high per-cell library construction costs which restrict the number of cells that can be assessed, thus imposing limitations on the ability to quantitatively measure genomic heterogeneity within a tissue. Here, we present Single cell Combinatorial Indexed Sequencing (SCI-seq) as a means of simultaneously generating thousands of low-pass single cell libraries for the purpose of somatic copy number variant detection. In total, we constructed libraries for 16,698 single cells from a combination of cultured cell lines, frontal cortex tissue from *Macaca mulatta*, and two human adenocarcinomas. This novel technology provides the opportunity for low-cost, deep characterization of somatic copy number variation in single cells, providing a foundational knowledge across both healthy and diseased tissues.

## Introduction

The booming field of single cell sequencing continues to shine light on the abundance and breadth of genomic heterogeneity between cells in a variety of contexts, including somatic gains or losses of megabasepair-sized regions of the genome in the mammalian brain^1–4^, and tumor heterogeneity and clonal evolution^5–7^. Single cell genome sequencing studies have taken one of two approaches: high depth of sequencing per cell for purposes of single nucleotide variant detection^2,8^, or low-pass sequencing to identify copy number variants (CNVs) and the presence of aneuploidy^1,9,10^. In the latter approach, the lack of a method to cost-effectively produce high numbers of single cell libraries has prevented the ability to quantitatively measure the frequency of CNV-harboring cells at population-level scale, or provide a robust analysis of heterogeneity in the context of cancer^11^.

Recently, we established a method to simultaneously produce thousands of individually barcoded libraries of linked sequence reads using a combinatorial indexing strategy with transposase-based library construction^12^ and applied the technology to haplotype resolution^13^ as well as contig scaffolding for *de novo* genome assembly^14^. The combinatorial indexing concept, which involves two tiers of library indexing (transposase stage and PCR stage) was integrated with the widely utilized assay for transposase accessible chromatin (ATAC-seq)^15^, to simultaneously produce profiles of active regulatory elements in thousands of single cells^16^ (scATAC-seq, **Fig. 1a**). In this method approximately 2,000 isolated nuclei are deposited into each well of a 96-well plate by either dilution or Fluorescence Activated Nuclei Sorting (FANS)^5^. These nuclei are then incubated with the transposase enzyme complexed with universal DNA adaptors that contain a unique index corresponding to each well of the plate. Due to the physical constraints of closed chromatin, the transposase is only able to incorporate indexed adaptors at regions of open chromatin^15^. The nuclei remain intact during this process, thus utilizing the nuclear scaffold itself as an individual reaction compartment that contains an indexed sequencing library representative of regions of open chromatin. Furthermore, the transposase complex remains tightly bound after adaptor incorporation^13,14^, which prevents individual library molecules from diffusing out of the nucleus. The set of 96 wells are then pooled and then 15-25 of these randomly indexed nuclei are deposited by a second round of FANS into each well of one or more new 96-well plates. The probability of any two nuclei within the new well being derived from the same origin well is low (6-11%)^16^. Each new well is then uniquely indexed by PCR followed by pooling all individual reactions, purification, and sequencing. Therefore, at the end of this process, each sequence read contains two indexes: Index 1 from the transposase plate, and Index 2 from the PCR plate. For each of the possible index combinations, approximately 20% contain true single cell libraries with the remaining index combinations unused. As proof of principle for this method, Cusanovich and colleagues produced over 15,000 single cell ATAC-seq profiles and demonstrated the ability to robustly separate a mix of two cell types by their accessible chromatin landscapes^16^. We reasoned that a similar combinatorial indexing strategy could be extended to single cell whole genome sequencing.

**Figure 1.**
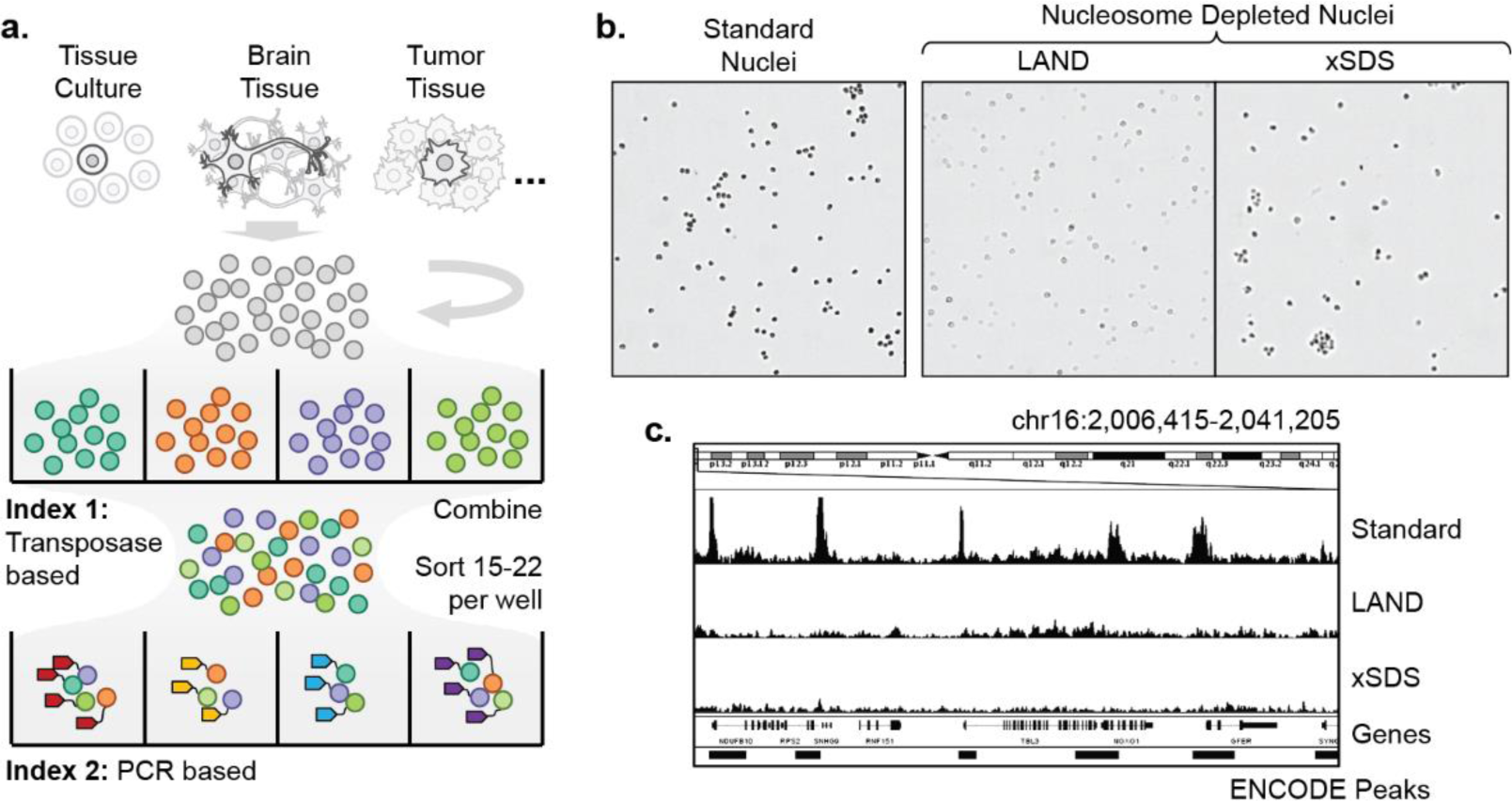
Single cell combinatorial indexing with nucleosome depletion. (**a**) Single cell combinatorial indexing workflow. (**b**) Standard isolated nuclei and nucleosome depleted nuclei using Lithium Assisted Nucleosome Depletion (LAND) or by crosslinking and SDS treatment (xSDS) all produce intact nuclear scaffolds. (**c**) Comparison of sequence coverage uniformity for bulk transposase-based library construction on standard or nucleosome depleted nuclei. Nucleosome depletion produces genome-wide uniform coverage that is not restricted to sites of chromatin accessibility.

Here we present our novel method, Single cell Combinatorial Indexing and Sequencing (SCI-seq), which harnesses the power of high throughput library construction and effectively removes the major bottleneck that is preventing deep characterizations of somatic aneuploidy. In order to achieve uniform coverage across the genome, as opposed to open chromatin enrichment from ATAC-seq, we developed two alternative strategies to deplete nucleosomes from bound DNA contained within the nuclear scaffold which is carried out prior to the combinatorial indexing library construction workflow. We demonstrate SCI-seq as a powerful method to produce thousands of low-pass whole genome sequencing libraries at low cost (<$1/cell) which can detect somatic aneuploidies and CNVs. We further show this method functions well across a variety of sources, including cultured cell lines, banked frozen Rhesus (Macaca mulatta) frontal cortex, and two human primary adenocarcinomas.

## Results

### Nucleosome depletion strategies geenrate uniform genome coverage

The key hurdle to adapt combinatorial indexing to produce uniformly distributed sequence reads of sufficient quantity for CNV detection is the removal of nucleosomes bound to genomic DNA without disturbing the nuclear scaffold. The scATAC-seq method is carried out on native chromatin in the nucleus, which only permits the conversion of DNA into viable library molecules within regions of open chromatin that encompass between 1-4% of the genome^17^. This restriction is desirable for epigenetic characterization; however, for purposes of CNV detection, it results in biological biases and severely limited read counts for each individual cell (median of roughly 3,000 unique reads per cell)^16^. We therefore developed two strategies: Lithium Assisted Nucleosome Depletion (LAND) and crosslinking with SDS treatment (xSDS), to unbind nucleosomes from genomic DNA while retaining nuclear scaffold integrity for SCI-seq library construction.

To test the viability of LAND and xSDS to deplete nucleosomes within nuclei, we first performed ATAC-seq preparations on bulk nuclei (30,000) from the HeLa S3 cell line for which chromatin accessibility and genome structure has been extensively profiled^18,19^. Libraries were constructed from standard isolated nuclei, nuclei exposed to Lithium diiodosalycylate (LAND), and nuclei that were crosslinked and subjected to SDS treatment in order to denature histones (xSDS). In all three cases, the nuclear scaffold remained intact – a key requirement for the SCI-seq workflow (**Fig. 1b**). Prepared nuclei were then carried through bulk sample ATAC-seq library construction^15^. The library prepared from standard nuclei produced the expected chromatin accessibility signal with a 10.8 fold enrichment of sequence reads aligning to annotated HeLa S3 accessibility sites. Both the LAND and xSDS preparations had substantially lower enrichment at 2.8 and 2.2 fold respectively, close to the expected distribution for a random shotgun sequencing library of 1.4 fold (**Fig. 1c**). Furthermore, the projected number of unique sequence reads present in the bulk sample LAND and xSDS preparations were 1.7 billion and 798 million respectively, much greater than for the standard ATAC-seq library at 170 million, indicating that a larger proportion of the genome was converted into viable sequencing molecules. Based on these results, we concluded that both LAND and xSDS methods are viable means of nucleosome depletion and facilitate a substantial increase in sequencing library complexity while retaining nuclear scaffold integrity.

### SCI-seq with nucleosome depletion

To assess the performance of nucleosome depletion with our single cell combinatorial indexing workflow, we chose to first focus on the euploid lymphoblastoid cell line GM12878, as it is in abundant supply and has been deeply profiled by a number of studies^13,14,18^. In order to test different Lithium diiodosalycylate concentrations and flow sorting conditions for the LAND method of nucleosome depletion, we generated seven SCI-seq libraries, six of which were successful with the seventh failing due to poor FANS gating on the second (post-transposition) sort. Each of these preparations were carried through using a single 96-well plate at the PCR indexing stage. For the xSDS method of nucleosome depletion we constructed a single SCI-seq library with 3×96-well plates at the PCR indexing stage to achieve high cell counts. To serve as a comparison to traditional single cell sequencing methods, we prepared 42 single cell libraries using quasi-random priming (QRP, 40 passing QC) and 51 single cell libraries using degenerate oligonucleotide primed PCR (DOP, 45 passing QC). Finally, we karyotyped an additional 50 cells to serve as a non-sequencing means of aneuploidy measurement.

For each SCI-seq preparation, the number of potential index combinations is 96 (transposase indexing) × N (PCR indexing wells, 96 per plate); however, not all of the index combinations represent a single cell library, as each PCR well only contains 15-25 transposase-indexed nuclei. To identify index combinations that contain a true single cell library, we first generated a log_10_ transformed histogram of unique (*i.e.* non-PCR duplicate), high-quality (MQ ≥ 10) aligned reads for each potential index combination. For high-quality preparations, this resulted in a bimodal distribution comprised of a low-read count, noise component taking up 1-3% of the total raw reads and centered between 50 and 200 reads, and a high-read count, single cell component made up of the remaining reads and centered between 10,000 and 100,000 reads (**Fig 2a,b**). We then identified the mean and standard deviation of each component using a mixed model and defined the minimum unique read count threshold for an index to be considered a single cell library. The threshold was determined to be either the greater of one standard deviation (in log_10_ space) below the mean of the single cell component or 100-fold greater than the mean of the noise component and at a minimum of 1,000 reads for libraries sequenced to very low depth. Using this approach, we identified a total of 4,643 single cell libraries across the six successful SCI-seq preparations that used LAND for nucleosome depletion and 3,123 from the single xSDS preparation.

**Figure 2.**
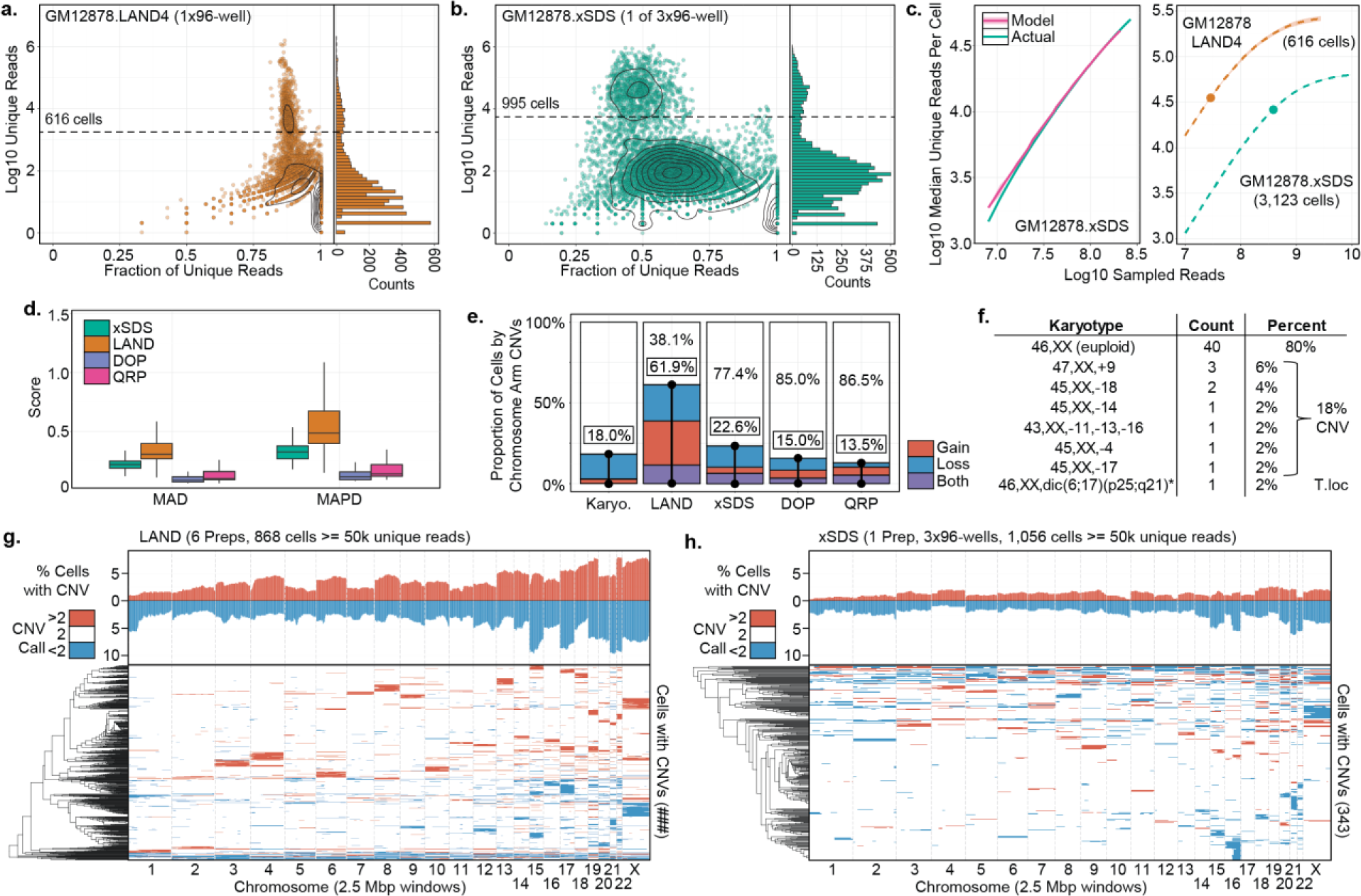
Comparison of LAND and xSDS nucleosome depletion methods with SCI-seq. (**a**) Log10 unique read count (y-axis) and histogram (right panel), by fraction of unique reads (x-axis) to indicate complexity for one of the seven SCI-seq preparations on GM12878 using LAND nucleosome depletion. Contour lines represent point density, dashed line represents single cell read cutoff. (**b**) As in (**a**) but for a SCI-seq preparation using xSDS for nucleosome depletion for one of the three PCR plates and sequenced to a greater depth than the LAND preparation. (**c**) Left, model built on downsampled reads for the GM12878 xSDS preparation and used to predict the full depth of coverage. Right, Projections for one of the LAND preparations and the full xSDS preparation. Points represent actual depth of sequencing. (**d**) Coverage uniformity scores for SCI-seq using LAND or xSDS for library construction and for two standard methods of single cell sequencing: quasi-random priming (QRP) and degenerate oligonucleotide PCR (DOP). (**e**) Summary of the percentage of cells showing aneuploidy at the chromosome arm level across all preparations. (**f**) Karyotyping results of 50 GM12878 cells. (**g-h**) Summary of windowed copy number calls and clustering of GM12878 single cells produced using LAND (**g**) or xSDS (**h**). Top represents a chromosome-arm scale summary of gain or loss frequency for all cells and bottom is the clustered profile for cells that contain at least one CNV call.

To confirm that the identified single cell index combinations are truly single cells, we carried out four SCI-seq library preparations on a mix of human (GM12878 or HeLa S3) and mouse (3T3) cells using LAND, and one preparation using xSDS for nucleosome depletion. Across the four LAND preparations we obtained 2,369 single cell libraries of which 2,248 (94.9%) had ≥ 90% of their reads that aligned solely to either the human or mouse genome. Similarly, the xSDS preparation produced 1,367 single cells with 1,242 (90.9%) having ≥ 90% of their reads aligning solely to one of the two species’ genomes. These experiments confirm that the single cell libraries we identified are indeed true single cells with index collision rates similar to scATACseq^16^ or high throughout single cell RNA-seq technologies^20^.

The total unique read count produced for each single cell library in a SCI-seq preparation is a function of library complexity (unique molecule count in library) and depth of sequencing. Due to the inhibitive cost of deeply sequencing *every* library preparation during method development, we implemented a nonlinear model to project the anticipated read count and PCR duplicate percentage that would be achieved with increased sequencing depth as a means of library quality assessment (**Fig. 2c**, Methods). These projections were used to calculate the expected median unique read count for cells in each library if it were sequenced to near saturation. We then identified the sequencing depth in which a median of 50% of the reads across cells are PCR duplicates (M50) which represents the point in which diminishing returns of further sequencing become excessive (*i.e.* greater than 50% of additional sequenced reads provide no additional information). We also identified the number of cells in each preparation that have the potential to produce sufficient read counts for low-resolution copy number calling, as well as a number of additional metrics. To evaluate our projections, we validated our approach on libraries sequenced to a higher depth (*e.g.* the xSDS preparation of GM12878) by building a model on a subset of the sequence reads and comparing the projected metrics with those from the actual high-depth sequencing. This analysis showed our model accurately predicted the mean and median read count within a median of 0.02% (maximum 2.25%, mean 0.41%) across all libraries.

Coverage uniformity was assessed using two previously described metrics – mean absolute deviation (MAD)^21^, and mean absolute pairwise deviation (MAPD)^2^, which indicated substantially increased uniformity using the xSDS nucleosome depletion strategy over the lithium-based LAND method (MAD: mean 1.57-fold improvement, *p* = <1x10^-15^; MAPD: 1.70-fold improvement, *p* = <1x10^-15^, Welch’s t-test), however the deviation of the xSDS preparation is still greater than for QRP and DOP methods, though similar to multiple displacement amplification methods of single cell sequencing (**Fig. 2d**)^21^. When further examining coverage, we observed a number of loci in which read pairs overlapped, with the vast majority overlapping at exactly 9 bp for both LAND and xSDS SCI-seq libraries, which is the length of DNA copied during the transposase incorporation reaction^22^, suggesting that we are sequencing library molecules on either side of the same transposase insertion event.

### Copy number variant calling using SCI-seq

Many of the SCI-seq libraries prepared during methods development were not sequenced deeply which resulted in a number of single cell libraries that lacked sufficient depth for accurate CNV calling. We therefore proceeded with cells for which a minimum of 50,000 unique, high quality aligned reads were produced (868 across LAND preparations, 1,056 for the xSDS preparation of GM12878 libraries), which we found to be sufficient for low-resolution analysis of CNVs. We then used the web-based tool, Ginkgo^21^, Circular Binary Segmentation (CBS)^23^, and a Hidden Markov Model (HMM)^24^, with variable-sized genomic windows with a target median size of 2.5 million basepairs (Mbp) for CNV calling. Prior to using the CBS or HMM approach, we first corrected for GC bias in our data on a per-read basis (Methods), as our only source of amplification bias is during PCR, and is therefore amplicon-specific. We then took a conservative approach where we retained only the intersected calls of the three methods for further analysis.

In order to compare our sequencing-based single cell CNV calls to the frequency of aneuploidy observed in the karyotyping analysis of GM12878, we included only those that spanned the majority (≥ 80%) of a chromosome arm to produce estimates of aneuploidy frequency at a comparable resolution (**Fig. 2e-g**). Consistent with the observed coverage uniformity differences, nucleosome depletion using LAND followed by SCI-seq produced a high aneuploidy rate (61.9%), suggesting an abundance of false positives due to lack of coverage uniformity. However, the xSDS nucleosome depletion strategy with SCI-seq resulted in an aneuploidy frequency of 22.6%, much closer to the karyotyping results (18% of cells harboring a chromosome-scale copy number alteration, **Fig. 2f**), and the two standard methods of single cell sequencing at 15.0% and 13.5% for DOP and QRP respectively.

We posited that the high variability in coverage uniformity observed using LAND may be due to cell cycle, with a proportion of cells in early- to mid- S-phase which might not have been excluded during flow sorting. However, after we arrested GM12878 cells in G0/G1 phase by serum starvation and proceeded through SCI-seq, the high variability persisted. Since the chromatin accessibility signal is essentially absent in libraries prepared from LAND nuclei, we believe that the variability may be due to DNA interactions with the nuclear scaffold itself, which are not removed by Lithium diiodosalycylate exposure^25^, and may warrant further investigation. Lastly, we prepared an additional three LAND nucleosome depletion SCI-seq libraries on HeLa S3 cells to characterize the performance in an aneuploid context which produced a total of 2,361 single cell libraries that largely conformed to the well-characterized HeLa copy number profile^19^.

While both LAND and xSDS methods of nucleosome depletion were able to produce SCI-seq libraries with genome-wide sequence coverage, each approach has its own advantages and disadvantages. Notably, LAND tends to produce higher read counts per cell (e.g. an M50 of 763,813 for one of the HeLa LAND preparations), albeit with increased bias of coverage. It may be possible to account for these biases if they are systematic, or to use other metrics (e.g. incidences of overlapping reads) for copy number calling. The bias may not hinder the interrogation of other properties, such as DNA methylation, where increased read counts would be preferred. xSDS typically results in fewer unique reads per single cell (M50 of 63,223 for the GM12878 preparation) but provides much more uniform coverage and allows for robust low-pass CNV calling on a high number of cells with frequencies in concordance with established methods and karyotyping analysis. Lastly, it may be possible to increase the read counts obtained from xSDS with SCI-seq by further optimizing the crosslinking and crosslink reversal processes.

### Copy number variation in the Rhesus brain

Estimates of the frequency of aneuploidy and large-scale copy number variation in the mammalian brain have varied widely, spanning a range of <5% to 33%^1–4^. This uncertainty largely stems from the inability to profile sufficient numbers of single cells to produce quantitative measurements. The Rhesus macaque is an ideal model for quantifying the abundance of aneuploidy in the brain, as human samples are challenging to acquire, with the majority of biopsies obtained from unhealthy individuals (e.g. epilepsy), and are further confounded by a high variability of lifetime environmental exposures that may influence mutation rate. Furthermore, the Rhesus brain is phylogenetically, structurally and physiologically more similar to humans than rodents^26^.

Without knowing the effects that freezing may have on nuclear scaffold integrity and the SCI-seq workflow, we carried out both LAND and xSDS nucleosome depletion methods in parallel along with two traditional methods of single cell sequencing: 38 cells using QRP (35 passing QC), and 35 cells using DOP (30 passing QC) all from adjacent sections of frontal cortex from the same individual (Individual 1). We prepared a low capacity (16 wells at the PCR indexing stage) SCI-seq library using LAND for nucleosome depletion from a ~150 mm^3^ sample which performed well, resulting in 340 single cells with a median unique read count of 141,449 and 248 cells with ≥ 50,000 unique reads. Our xSDS preparation from the adjacent ~150 mm^3^ sample generated 171 single cell libraries with a median unique read count of 55,142 and 92 cells passing the 50,000 read count threshold. The number of cells produced using SCI-seq with xSDS was lower than expected when compared to cell culture experiments largely due to challenges in the nuclei isolation and FANS. We believe that increasing the amount of tissue entering the workflow may produce sufficient numbers of nuclei for more selective gating during the flow sorting process which should translate to increased numbers of passing single cell libraries.

Across all methods of library construction we observed greater discrepancies between the three CNV calling approaches than in the human analyses (**Supplementary Fig. 9-12**). We believe that this variability is due to the lower quality of the Rhesus reference genome when compared to human, emphasizing the need for so-called “platinum” quality reference genomes^27^. These discrepancies included difficulty in identifying baseline ploidy for Ginkgo (a step exclusive to that method), a bias towards calling only large CNVs by CBS, and the HMM approach producing a greater number of calls, though possibly at the cost of a higher false positive rate. Interestingly, the challenges with Ginkgo were only present for QRP and DOP and not in either of the SCI-seq preparations. We therefore focused on the HMM results for sub-chromosomal calls (**Fig. 3a,b**) and the intersection of CBC and HMM calls for aneuploidy analysis, retaining only calls that span ≥ 80% of a chromosome (**Fig. 3c**).

**Figure 3.**
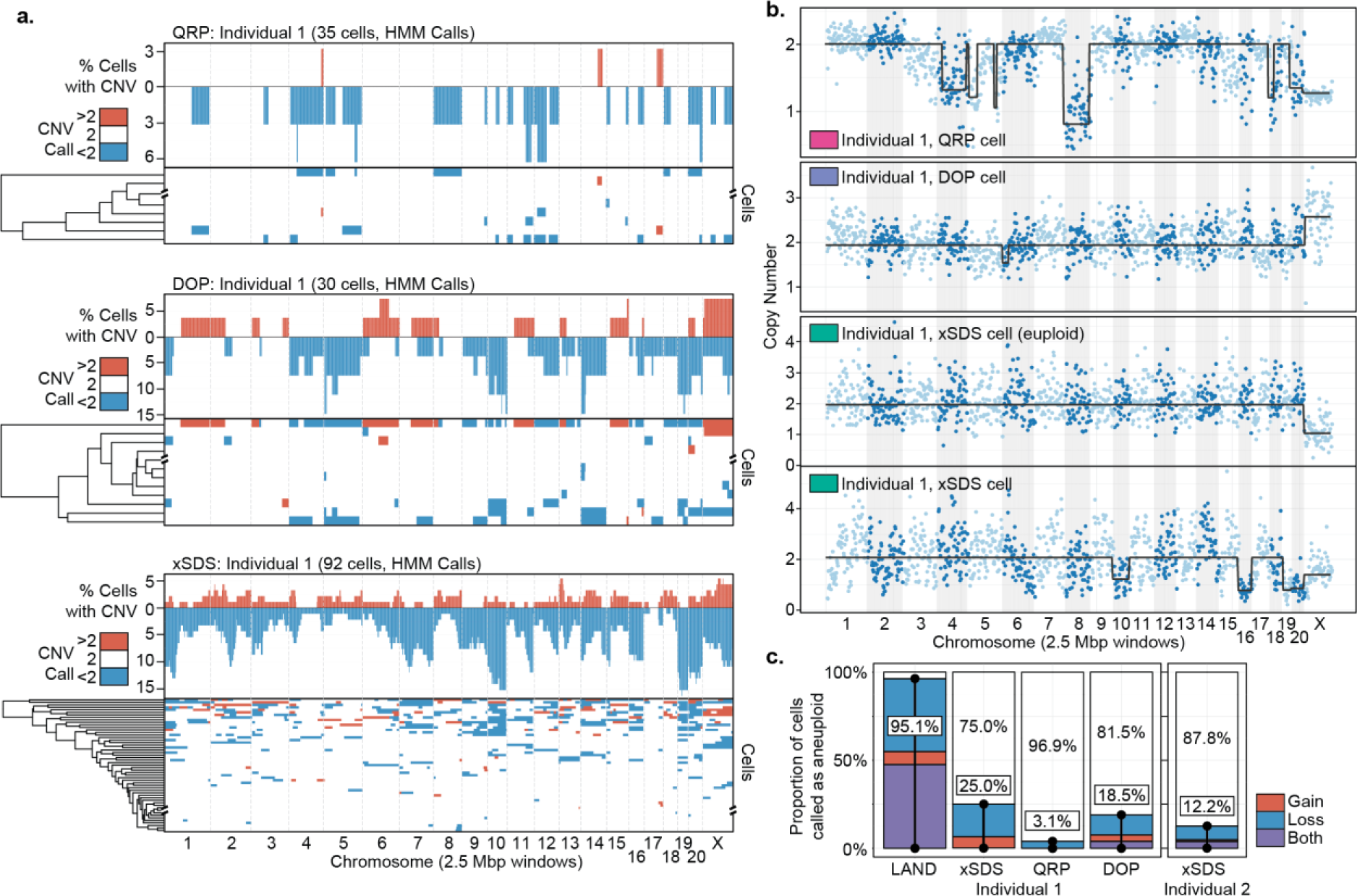
Somatic CNVs in the Rhesus brain. (**a**) Summary of CNV calls using the two standard methods of library construction (Quasi-Random Priming, QRP; and Degenerate Oligonucleotide Primed PCR, DOP) and SCI-seq using xSDS for nucleosome depletion on adjacent sections of Rhesus frontal cortex. Depicted calls were made using a Hidden Markov Model method (HMM). The top protion represents a summary of CNV calls with the bottom portion showing clustered aneuploid cells with all fully-euploid cells collapsed at the axis break. (**b**) Example single cells with copy number variants, and one representative euploid cell for the SCI-seq preparation (HMM). (**c**) Frequency of aneuploidy as determined by each of the methods (HMM & CBS intersection, calls ≥ 80% of a chromosome).

Consistent with our cell line results, the LAND preparation produced a much higher aneuploidy call rate (95.1%), suggestive of false positives stemming from coverage nonuniformity. Interestingly, the excess aneuploidy calls are highly enriched for the gain or loss of specific chromosomes (10, 16, 19, and 20), which were also more frequently called as aneuploid for the xSDS and DOP libraries. This is suggestive of a systematic bias that may be ameliorated by improving the reference assembly. The xSDS SCI-seq aneuploidy rate (25.0% of cells) was close to the DOP preparation (18.5%), with QRP producing a much lower rate (3.1%; **Fig. 3c**). Using xSDS, we prepared an additional library on a 200 mm^3^ section of frontal cortex from a second individual which produced 381 single cell libraries with a median depth of 62,731 and 213 libraries sequenced to at least 50,000 unique reads. Aneuploidy analysis of this second individual produced a rate similar to that of the first for both xSDS SCI-seq and DOP preparations at 12.1% of cells harboring a copy number variant chromosome, suggestive of low inter-individual variability, though a much larger cohort must be examined to provide definitive results.

### SCI-seq on primary tumor samples reveals clonal populations

One of the primary applications of single cell genome sequencing is in the profiling of tumor heterogeneity and understanding clonal evolution in cancer as it relates to treatment resistance^5–7^. We posited that the high throughput of SCI-seq and coverage uniformity using the xSDS method of nucleosome depletion may facilitate an in depth analysis of tumor heterogeneity by single cell CNV analysis. We therefore carried out xSDS SCI-seq on a freshly acquired stage III pancreatic ductal adenocarcinoma (PDAC) specimen measuring approximately 250 mm^3^. Our preparation resulted in 1,715 single cell libraries which were sequenced to a median unique read count of 49,272 per cell (M50 of 71,378) with 846 cells having at least 50,000 unique reads at the depth the library was sequenced and carried through CNV calling (**Fig. 4a**).

**Figure 4.**
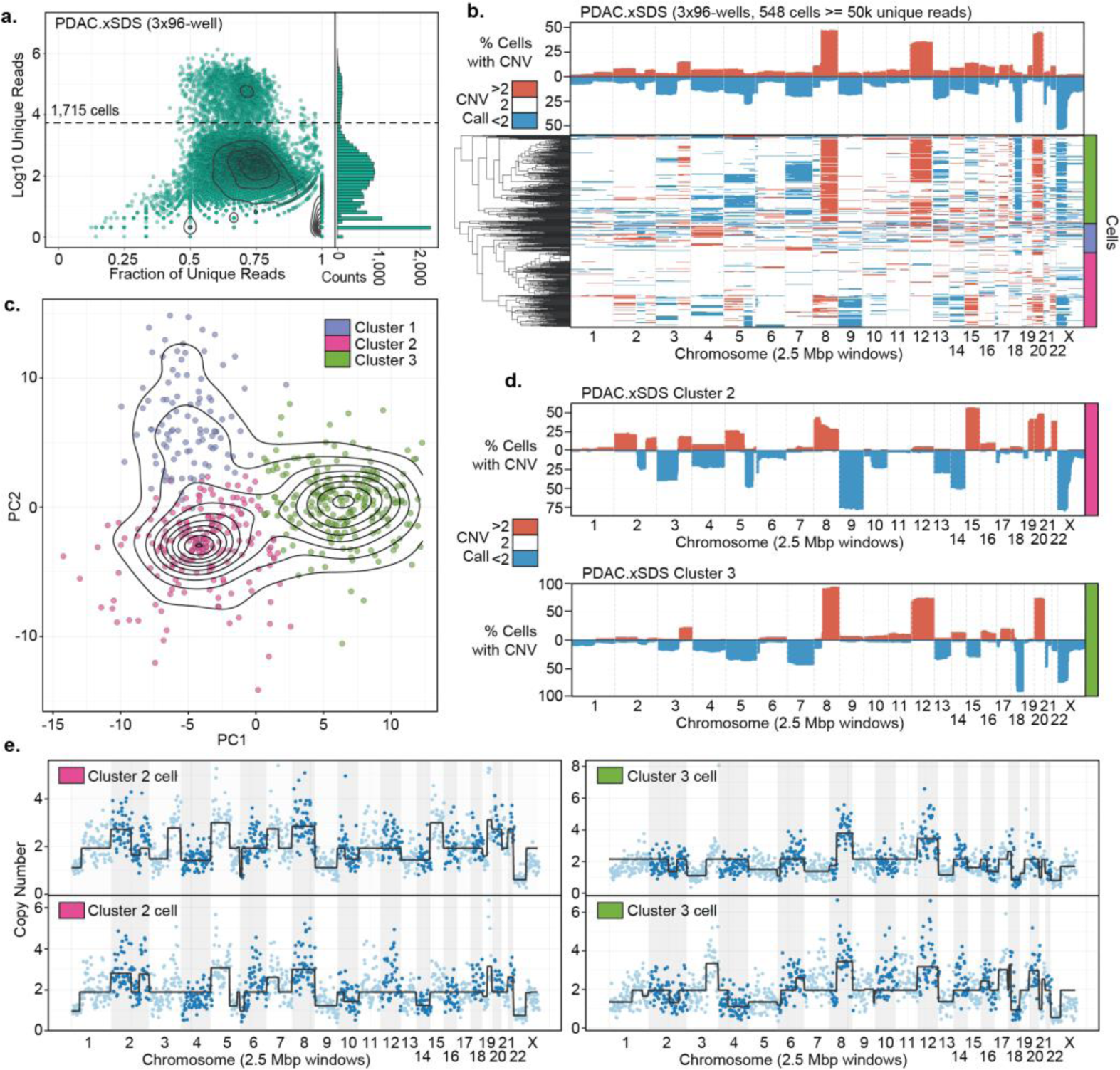
SCI-seq analysis of a stage III human Pancreatic Ductal Adenocarcinoma (PDAC). (**a**) SCI-seq library complexity as in Figure 1a. (**b**) CNV profile on non-euploid cells for the intersection of Ginkgo, CBS, and HMM call sets. (**c**) Principle component analysis and k-means clustering on CNV profiles. (**d**) CNV summaries of two distinct aneuploid subpopulations. (**e**) Representative single cell plots of CNV calls.

For both CBS and HMM copy number calling platforms, a baseline reference of expected euploid sequence coverage must be used, something that is not required for the Ginkgo platform. For analyses where the assumption that the vast majority of cells should be euploid (*i.e.* GM12878 and Rhesus experiments), the combined pool of cells can be used for this purpose; however this assumption does not hold true for a tumor sample for which a significant number of aneuploid cells are expected. We therefore first used the pooled reads from our GM12878 xSDS SCI-seq preparation to act as the coverage baseline which allowed us to identify a population of cells with no copy number variation at high confidence (298, 35.2%) to provide a new baseline specific to the individual (*i.e.* accounts for germline variation) and to the specific library preparation to carry out CNV calling on the remaining 548 cells.

The resulting CNV profile (**Fig. 4b**) revealed three distinct populations that were separated by performing a principle components analysis (PCA, **Fig. 4c**) on the copy number call matrix followed by k-means clustering. This produced one population that was comprised of predominantly euploid cells that were not removed during the baselining (99 cells, 18.1%), and then two tumor cell populations: cluster 1 comprised of 217 cells (39.6%) and cluster 2 comprised of 233 (42.5%) cells (**Fig 4d,e**). The cluster 2 population further split into two additional populations that were clearly identified in the HMM calls; however, the distinction was less clear in the intersection calls. Taken together, we were able to estimate an epithelial tumor cell purity of 53.2%, which is in line with the expected range for PDAC^28^, with the predominantly euploid population most likely comprised of a mix normal epithelial cells as well as infiltrating immune cells and vessels. Interestingly, the two subpopulations have significantly different CNV profiles, with the majority cells in cluster 2 having a deletion of chromosomes 9 and 14, and amplifications of chromosomes 15, 19q, and 22, all of which are not observed in cells assigned to cluster 3. Conversely, cluster 3 cells harbor a chromosome 12 amplification and an 18q deletion that is not observed in cluster 2. Both subpopulations exhibit a chromosome Xp deletion, and chromosome 8 amplification, though in cluster 3, this amplification is largely restricted to the q arm. We also observed a number of copy number calls that were only present in a subset of cells within a population. While a number of these represent subclonal variation, the highly conservative CNV calling approach we have taken which includes the intersect of three strategies, likely under-calls some of these variants which would imply they are closer to fixation in any given clone, as observed in the profiles for each individual calling method.

We next applied SCI-seq to a frozen stage II rectal cancer tissue sample measuring approximately 500 mm^3^ using the xSDS workflow. Upon homogenization and lysis we noticed a high abundance of nuclear debris and ruptured nuclei which resulted in a substantially decreased yield during the library construction process for which only 16 wells (as opposed to ≥ 96) of PCR-stage indexing were able to be carried out. This produced 146 single cell libraries that were sequenced to a median single cell unique read count of 71,378 (M50 of 352,168) and 111 cells sequenced to a depth of at least 50,000. As with the PDAC sample, we were able to identify a largely euploid population (54 cells, 48.6%) which was then used to baseline the CNV calling for the remaining aneuploid cells (57 cells, 51.4%). However, unlike the PDAC preparation, we did not observe clear clonal populations with a number of sporadic calls. This could be a result of irradiation, a common treatment for rectal cancers, or that the nuclear integrity was compromised during tissue homogenization, underscoring the challenge of producing high-quality single cell or nuclei suspensions shared by all single cell methods^11^.

## Discussion

The inability to cost-effectively produce single cell genome sequencing libraries in the numbers necessary for quantitative measurements has been the crux of somatic aneuploidy research. Here we have developed a novel approach, SCI-seq, to combinatorially scale the production of low-pass single cell genome sequencing libraries for aneuploidy and copy number variant detection. To achieve this, we implemented two nucleosome depletion strategies, LAND and xSDS, each of which have their respective advantages and disadvantages. We found that LAND is able to produce a greater number of reads per cell but at the cost of decreased coverage uniformity, whereas xSDS produces less reads with improved coverage uniformity. The reduced read counts of xSDS is most likely due to inefficient crosslink reversal, and optimization of this process or the exploration of alternative crosslinking agents may result in substantially increased read counts. We conclude that xSDS is the preferred SCI-seq nucleosome depletion method for CNV analysis while LAND may be preferred for adapting the SCI-seq platform for the interrogation of other genomic or epigenomic properties where uniformity is less essential.

One of the major limitations of SCI-seq is that, without pre-amplification of genomic DNA, there is a theoretical maximum coverage of one read at any given position per copy present. This prevents the possibility of complete or deep coverage that is necessary for single nucleotide variant detection. A future development could include *in situ* pre-amplification within the intact nuclear scaffold after nucleosome depletion may be possible; however amplification will have to be tightly controlled to prevent nuclei from rupturing and may be limited to cell types with larger nuclei. Another potential strategy would be to integrate SCI-seq with T4 *in vitro* transcription, as has been demonstrated with THS-seq^29^, an ATAC-seq alternative, to boost the resulting coverage. Lastly, SCI-seq in its current form is only compatible with cell types in which a nuclear scaffold exists, and therefore cannot be performed on prokaryotic cells. It has been shown that crosslinking of genomic DNA within bacteria can produce an enrichment of proximal reads^30,31^, which may be sufficient for genome compartmentalization of individual cells; however, the additional challenge of flow sorting the resulting aggregates may prove difficult.

We believe that SCI-seq represents a new high throughput platform to accurately profile somatic heterogeneity or tumor evolution. In total we produced 16,698 single cell libraries (of which 5,395 were sequenced to a depth sufficient for CNV calling) from myriad samples using SCI-seq, including primary tissue isolates representative of the two major areas of single cell genome research: aneuploidy in the brain, and in cancer. In addition to the advantages of throughput, the platform does not require specialized microfluidics equipment or droplet emulsification techniques, and it can be performed on isolated nuclei, thus allowing it to be carried out on tissue in which cells are difficult to dissociate. While further optimization is possible, as with any new method, we believe that the throughput provided by SCI-seq will open the door to deep quantification of mammalian somatic genome stability as well as serve as a platform to assess other properties of single cells including DNA methylation and chromatin architecture.

## Methods

### Sample preparation and nuclei isolation

Tissue culture cell lines were trypsinized then pelleted if adherent (HeLa S3, ATCC CCL-2.2; NIH/3T3, ATCC CRL-1658) or pelleted if grown in suspension (GM12878, Coriell; karyotyped at the OHSU Research Cytogenetics Laboratory), followed by one wash with ice cold PBS. They were then carried through crosslinking (for the xSDS method) or directly into nuclei preparation using Nuclei Isolation Buffer (NIB, 10mM TrisHCl pH7.4, 10MM NaCl, 3mM MgCl2, 0.1% igepal, 1x protease inhibitors (Roche, Cat. 11873580001)) with or without nucleosome depletion. Tissue samples (RhesusFcx1, RhesusFcx2, PDAC, CRC) were dounce homogenized in NIB then passed through a 35µm cell strainer prior to nucleosome depletion. The frozen Rhesus frontal cortex samples, RhesusFcx1 (4 yr. male) and RhesusFcx2 (9 yr. male), were obtained from the Oregon National Primate Research Center as a part of their aging nonhuman primate resource.

### Standard Single Cell Library Construction

Single cell libraries constructed using quasi-random priming (QRP) and degenerate oligonucleotide primed PCR (DOP) were prepared from isolated nuclei without nucleosome depletion and brought up to 1 mL of NIB, stained with 5 μL of 5mg/ml DAPI (Thermo Fisher, Cat. D1306) then FANS sorted on a Sony SH800 in single cell mode. One nucleus was deposited into each single well containing the respective sample buffers. QRP libraries were prepared using the PicoPlex DNA-seq Kit (Rubicon Genomics, Cat. R300381) according to the manufacturer’s protocol and using the indexed PCR primers provided in the kit. DOP libraries were prepared using the SeqPlex DNA Amplification Kit (Sigma, Cat. SEQXE-50RXN) according to the manufacturer’s protocol, but with the use of our own custom PCR indexing primers that contain 10 bp index sequences. To avoid over-amplification, all QRP and DOP libraries were amplified with the addition of 0.5 µL of 100X SYBR Green (FMC BioProducts, Cat. 50513) on a BioRad CFX thermocycler in order to monitor the amplification and pull reactions that have reached mid-exponential amplification.

### Nucleosome Depletion

*Lithium assisted nucleosome depletion (LAND)*: Prepared Nuclei were pelleted and resuspended in NIB supplemented with 200 μL of 12.5 mM lithium 3,5-diiodosalicylic acid (referred to as Lithium diiodosalycylate in main text, Sigma, Cat. D3635) for 5 minutes on ice prior to the addition of 800 μL NIB and then taken directly into flow sorting.

*Crosslinking and SDS nucleosome depletion (xSDS)*: Crosslinking was achieved by incubating cells in 10 mL of media (cell culture) or nuclei in 10 mL of HEPES NIB (20mM HEPES, 10MM NaCl, 3mM MgCl2, 0.1% igepal, 1x protease inhibitors (Roche, Cat. 11873580001)) (tissue samples) containing 1.5% formaldehyde at room for 10 minutes. The crosslinking reaction was neutralized by bringing the reaction to 200mM Glycine (Sigma, Cat. G8898-500G) and incubating on ice for 5 minutes. Cell culture samples were crosslinked and then washed once with 10 ml ice cold 1x PBS and had nuclei isolated by incubating in NIB buffer on ice for 20 minutes and pelleted once again. Nuclei were then resuspended in 800 uL 1x NEBuffer 2.1 (NEB, Cat. B7202S) with 0.3% SDS(Sigma, Cat. L3771) and incubated at 42°C with vigorous shaking for 30 minutes in a thermomixer (Eppendorf). SDS was then quenched by the addition of 200 µL of 10% Triton-X100 (Sigma, Cat. 9002-93-1) and incubated at 42°C with vigorous shaking for 30 minutes.

### Combinatorial indexing via tagmentation and PCR

Nuclei were stained with 5 μL of 5mg/ml DAPI (Thermo Fisher, Cat. D1306) and passed through a 35µm cell strainer. A 96 well plate was prepared with 10 μL of 1x Nextera^®^ Tagment DNA (TD) buffer from the Nextera^®^ DNA Sample Preparation Kit (Illumina, Cat. FC-121-1031) diluted with NIB in each well. A Sony SH800 flow sorter was used to sort 2,000 single nuclei into each well of the 96 well tagmentation plate in fast sort mode. Next, 1 μL of a uniquely indexed 2.5 µM transposase-adaptor complex (transposome) was added to each well. These complexes and associated sequences are described in Amini *et. al.* 2015, Ref. 13. Reactions were incubated at 55°C for 15 minutes. After cooling to room temperature, all wells were pooled and stained with DAPI as previously described. A second 96 well plate, or set of 96 well plates, were prepared with each well containing 8.5 μL of a 0.058% SDS, 8.9 nM BSA solution and 2.5 μL of 2 uniquely barcoded primers at 10 µM. 22 post-tagmentation nuclei from the pool of 96 reactions were then flow sorted on the same instrument but in single cell sort mode into each well of the second plate and then incubated in the SDS solution at 55°C for 5 minutes to disrupt the nuclear scaffold and disassociate the transposase enzyme. Crosslinks were reversed by incubating at 68°C for an hour (xSDS). SDS was then diluted by the addition of 7.5 µL of Nextera^®^ PCR Master mix (Illumina, Cat. FC-121-1031) as well as 0.5 µL of 100X SYBR Green (FMC BioProducts, Cat. 50513) and 4 µL of water. Real time PCR was then performed on a BioRad CFX thermocycler by first incubating reactions at 72°C for 5 minutes, prior to 3 minutes at 98°C and 15-20 cycles of [20 sec. at 98°C, 15 sec. at 63°C, and 25 sec. at 72°C]. Reactions were monitored and stopped once exponential amplification was observed in a majority of wells. 5 µL of each well was then pooled and purified using a Qiaquick PCR Purification column (Qiagen, Cat. 28104) and eluted in 30 µL of EB.

### Library quantification and sequencing

Libraries were quantified between the range of 200bp and 1kb on a High Sensitivity Bioanalyzer kit (Agilent, Cat. 5067-4626). Libraries were sequenced on an Illumina NextSeq^®^ 500 loaded at 0.8 pM with a custom sequencing chemistry protocol (Read 1: 50 imaged cycles; Index Read 1: 8 imaged cycles, 27 dark cycles, 10 imaged cycles; Index Read 2: 8 imaged cycles, 21 dark cycles, 10 imaged cycles; Read 2: 50 imaged cycles) using custom sequencing primers described in Amini *et. al.* 2015, Ref. 13. QRP and DOP libraries were sequenced using standard primers on the NextSeq^®^ 500 using high-capacity 75 cycle kits with dual-indexing. For QRP there is an additional challenge that the first 15 bp of the read are highly enriched for “G” bases, which are non-fluorescent with the NextSeq^®^ 2-color chemistry and therefore cluster identification on the instrument fails. We therefore sequenced the libraries using a custom sequencing protocol that skips this region (Read 1: 15 dark cycles, 50 imaged cycles; Index Read 1: 10 imaged cycles; Index Read 2: 10 imaged cycles).

### Sequence Read Processing

Sequence runs were processed using bcl2fastq (Illumina Inc., version 2.15.0) with the --create-fastq-for-index-reads and --with-failed-reads options to produce fastq files. Index reads were concatenated (36 bp total) and used as the read name with a unique read number appended to the end. These indexes were then matched to the corresponding index reference sets allowing for a hamming distance of two for each of the four index components (i7-Transposase (8 bp), i7-PCR (10 bp), i5-Transposase (8 bp), and i5-PCR (10bp)), reads matching a quad-index combination were then renamed to the exact index (and retained the unique read number) which was subsequently used as the cell identifier. Reads were then adaptor trimmed, then paired and unpaired reads were aligned to reference genomes by Bowtie2^32^ and merged. Human preparations were aligned to GRCh37, Rhesus preparations were aligned to RheMac8, and Human/Mouse mix preparations were aligned to a combined human (GRCh37) and mouse (mm10) reference. Aligned bam files were subjected to PCR duplicate removal using a custom script that removes reads with identical alignment coordinates on a per-barcode basis along with reads with an alignment score less than 10 as reported by Bowtie2.

### Single Cell Discrimination

For each PCR plate, a total of 9,216 unique index combinations are possible (12 i7-Transposase indexes × 8 i5-Transposase indexes × 12 i7-PCR indexes × 8 i5-PCR indexes), for which only a minority should have a substantial read count, as the majority of index combinations should be absent – *i.e.* transposase index combinations of nuclei that were not sorted into a given PCR well. These “empty” indexes typically contain very few reads (1-3% of a run) with the majority of reads falling into *bona fide* single cell index combinations (97-99% of a run). The resulting histogram of log_10_ unique read counts for index combinations (**Supplementary Fig. N**) produces a mix of two normal distributions: a noise component and a single cell component. We then used the R package “mixtools” to fit a mixed model (normalmixEM()) to identify the proportion (λ) mean (µ) and standard deviation (σ) of each component. The read count threshold to qualify as a single cell library was taken to be the greater of either one standard deviation below the mean of the single cell component in log_10_ space, or 100 fold greater than the mean of the noise component (+2 in log_10_ space), and had to be a minimum of 1,000 unique reads.

### Human-Mouse Mix Experiments

We took one of two approaches to mix human and mouse cells: i) mixing at the cell stage (HumMus.LAND1 and HumMus.LAND2) or ii) mixing at the nuclei stage (HumMus.LAND3, HumMus.LAND4, and HumMus.xSDS). The reason we employed the latter was to control for nuclei crosslinking or agglomerating together that could result in doublets. Libraries were constructed as described above, for instances where two distinct DAPI-positive populations were observed during flow sorting, included both populations in the same gate so as not to skew proportions. Reads were processed as in other experiments, except reads were instead aligned to a reference comprised of GRCh37 (hg19) and mm10. The mapping quality 10 filter effectively removed reads that aligned to conserved regions in both genomes and then for each identified single cell, reads to each species were tallied and used to estimate collision frequency.

### Library Depth Projections

To estimate the performance of a library pool if, or when, it was sequenced to a greater depth, we incrementally sampled random reads from each SCI-seq preparation across all index combinations including unaligned and low quality reads without replacement at every one percent of the total raw reads. For each point we identified the total number reads that are aligned with high quality (MQ ≥ 10) assigned to each single cell index and the fraction of those reads that are unique, non-PCR duplicates, as well as the corresponding fraction of total reads sampled that were assigned to that index. Using these points we fit both a nonlinear model and a Hanes-Woolfe transformed model to predict additional sequencing for each individual single cell library within the pool and projected out to a median unique read percentage across cells of 5%. To determine the accuracy of the models, we determined the number of downsampled raw reads of each library that would reach the point in which the median unique read percentage per cell was 90%, which is somewhat less than what was achieved for libraries that were sequenced at low coverage. We then subsampled the pre-determined number of reads for 30 iterations and built a new model for each cell at each iteration and then predicted the unique read counts for each cell out to the true sequencing depth that was achieved. The standard deviation of the true read count across all iterations for all cells was then calculated.

### Genome Windowing

Genomic windows were determined on a per-library basis using custom tools. For each chromosome the size of the entire chromosome was divided by the target window size to produce the number of windows per chromosome. The total read count for the chromosome summarized over the pool of all single cells (GM12878 for all human samples where absolute copy number was determined, as well as for each pooled sample where amplifications or deletions relative to the mean copy number were determined) was then divided by the window count to determine the mean read count per window. The chromosome was then walked and aligned reads from the pool tallied and a window break was made once the target read count per window was reached. Windows at chromosome boundaries were only included if they contained more than 75% of the average reads per window limit for that chromosome. By using dynamic windows we accounted for biases, such as highly repetitive regions, centromeres and other complex regions that can lead to read dropout in the case of fixed size bins^21^.

### GC Bias Correction

Reads were placed into the variable sized bins and GC corrected based on individual read GC content instead of the GC content of the dynamic windows. We posit that the large bin sizes needed for single cell analysis average out smaller scale GC content changes. Furthermore, SCI-seq does not involve pre-amplification where large regions of the genome are amplified, therefore GC bias originates solely from the PCR and is amplicon-specific. To calculate correction weights for the reads we compared the fraction of all reads with a given GC to the fraction of total simulated reads with the average insert size at the same GC fraction. This weight was then used in lieu of read counts and summed across all reads in a given window. All regions present in DAC blacklisted regions were excluded from analysis for the human sample analyses (http://genome.ucsc.edu/cgi-bin/hgFileUi?db=hg19&g=wgEncodeMapability)^18^. Following GC correction, all reads were normalized by the average number of reads per bin across the genome. Finally for each window we took the normalized read count of each cell and divided it by the pooled sample baseline to produce a ratio score.

### Measures of data variation

To measure data quality, we calculated three different measures of coverage dispersion: the median absolute deviation (MAD), the median absolute pairwise difference (MAPD) and the dispersion of reads. For the MAD and MAPD scores we calculated the median of the absolute values of all pairwise differences between neighboring bins that have been normalized by the mean bin count within the cell (log2 normalized ratios for the MAPD scores). For the dispersion we calculated the ratio of the variance and the mean value of all normalized bin values in a cell. The MAD and MAPD scores measure the dispersion of normalized binned reads due to technical noise, rather than due copy number state changes, which are less frequent^2,21^.

### Copy Number Variant Calling

CNV calling was performed on the windowed, GC corrected and bulk sample normalized reads with two available R packages that employ two different segmentation strategies: a Hidden Markov Model approach (HMMcopy, version 3.3.0, Ref. 24) and Circular Binary Segmentation (DNAcopy, version 1.44.0, Ref. 23). Values were Log2 transformed for input (2*log2 for CBS) and copy number calls were made based on the optimized parameters from Knouse et al. 2016, Ref. 10. For optimal sensitivity and specificity to detect copy number calls with sizes ≥5Mb we set the probability of segment extension (E) to 0.995 for HMM and for CBS we chose the significance level to accept a copy number change (α) to be 0.0001. The Log2 cutoffs for calling losses or gains were 0.4 and −0.35 for HMM and 1.32 and 0.6 for CBS. As an additional tool for CNV calling we used Ginkgo^21^, which uses an alternative method for data normalization. We uploaded bed files for each cell and a bulk down sampled bed file, which we created with Picard Tools (we used a down sample probability of 0.1). For the analysis we chose to segment single cells with the down sampled bulk bed file and when ploidy was known for the samples we created FACS files to force Ginkgo to normalize to that ploidy. Calls for the three methods were intersected either on a per-window basis or were filtered to only include calls that span ≥ 80% of a chromosome arm and then intersected for aneuploidy analysis. Principle components analysis of the intersected, window CNV calls for the PDAC sample was performed using R prcomp().

## Data Access

GM12878 and Rhesus sequence data are undergoing submission to the NCBI Sequence Read Archive (SRA) for unrestricted access. HeLa sequence data are undergoing submission to the database of Genotypes and Phenotypes (dbGaP), as a substudy under accession number phs000640. Human tumor samples are undergoing submission to dbGaP and are awaiting study accession assignment. All methods for making the transposase complexes are described in (Ref. 13).

## Acknowledgements

The genome sequence described/used in this research was derived from a HeLa cell line. Henrietta Lacks, and the HeLa cell line that was established from her tumor cells without her knowledge or consent in 1951, have made significant contributions to scientific progress and advances in human health. We are grateful to Henrietta Lacks, now deceased, and to her surviving family members for their contributions to biomedical research. The data generated from this research were submitted to the database of Genotypes and Phenotypes (dbGaP), as a substudy under accession number phs000640.

We thank the aging nonhuman primate resource at the Oregon National Primate Research Center for the banked Rhesus samples, the Brenden-Colson Center for Pancreatic Care for the pancreatic ductal adenocarcinoma sample, and the Knight Tissue Bank for the rectal adenocarcinoma sample. We thank Jay Shendure and Shendure lab members Darren Cusanovich, and Riza Daza for helpful advice and comments, and Martin Kircher for providing PCR-stage index sequences. We also thank Brian J. O’Roak for helpful discussions and manuscript suggestions. A.A. is supported by an Oregon Medical Research Foundation New Investigator Award. J.L.R. is supported by the Collins Medical Trust Foundation and Glenn/AFAR Scholarship for Research in the Biology of Aging. L. Carbone is supported by the Office of the Director/Office of Research Infrastructure Programs (OD/ORIP) of the NIH (grant no. OD011092).

## Author Contributions

A.A. designed and supervised all aspects of the study. A.A., S.A.V., and K.A.T. wrote the manuscript. All authors contributed and edited the manuscript. S.A.V. carried out all SCI-seq and GM12878 DOP library preparations, designed experiments, and performed all sequencing. A.A. and K.A.T. processed all sequence data and analyzed data. K.A.T. performed all copy number calling. J.L.R. constructed QRP and DOP libraries on Rhesus samples. A.J.F. prepared all GM12878 QRP library construction and co-prepared all SCI-seq libraries using xSDS for nucleosome depletion. M.H.W. provided tumor samples and aided in the analyses of those samples. L. Carbone supervised and provided all samples for Rhesus work. F.J.S. contributed to experimental design and contributed to the manuscript, L. Christiansen produced all transposase complexes used in this study.

